# Mutational burden of hepatoblastomas: a role for the *CX3CL1/CX3CR1* chemokine signaling pathway

**DOI:** 10.1101/555466

**Authors:** Talita Ferreira Marques Aguiar, Maria Prates Rivas, Silvia Costa, Tatiane Rodrigues, Juliana Sobral de Barros, Anne Caroline Barbosa, Mariana Maschietto, Renan Valieris, Gustavo Ribeiro Fernandes, Monica Cypriano, Silvia Regina Caminada de Toledo, Angela Major, Israel Tojal, Maria Lúcia de Pinho Apezzato, Dirce Maria Carraro, Carla Rosenberg, Cecilia Maria Lima da Costa, Isabela Werneck da Cunha, Stephen Frederick Sarabia, Dolores-López Terrada, Ana Cristina Victorino Krepischi

## Abstract

**Background:** Hepatoblastoma is an embryonal liver tumor supposed to arise from the impairment of hepatocyte differentiation during embryogenesis. CTNNB1 is the only recurrently mutated gene, and this relative paucity of somatic mutations poses a challenge to risk stratification and development of targeted therapies.

**Methods:** In this study, we investigated by exome sequencing the burden of somatic mutations in a cohort of 10 hepatoblastomas, including a congenital case.

**Results:** Our data disclosed a low mutational background with only three recurrently mutated genes: CTNNB1 and two novel candidates, CX3CL1 and CEP164. The major finding was a recurrent mutation (A235G) identified in two hepatoblastomas at the CX3CL1 gene; evaluation of RNA and protein expression revealed up-regulation of CX3CL1 in tumors. The analysis was replicated in two independents cohorts, substantiating that an activation of the CX3CL1/CX3CR1 pathway occurs in hepatoblastomas, with a predominance of these proteins in the cytoplasm of tumor cells. These proteins were not detected in the infiltrated lymphocytes of inflammatory regions of the tumors, in which they should be expressed in normal conditions, whereas necrotic regions exhibited negative tumor cells, but strongly positive infiltrated lymphocytes. Our data suggested that CX3CL1/CX3CR1 upregulation may be a common feature of hepatoblastomas, potentially related to chemotherapy response and progression. In addition, three mutational signatures were identified in hepatoblastomas, two of them with predominance of either the COSMIC signatures 1 and 6, found in all cancer types, or the COSMIC signature 29, related only with tobacco chewing habit; a third novel mutational signature presented an unspecific pattern with an increase of C>A mutations.

**Conclusions:** Overall, we present here evidence that CX3CL1/CX3CR1 chemokine signaling pathway is likely involved with hepatoblastoma tumorigenesis or progression, besides reporting a novel mutational signature specific to a hepatoblastoma subset.

## INTRODUCTION

Hepatoblastoma (HB) is the most common malignant liver tumor in the pediatric population [1], supposedly derived from hepatocyte precursors [2]. Although rare, there is a trend towards an increasing prevalence of HBs over the last years [3]. The cause of this rising in incidence is still unknown, but a possible explanation would be the increasing survival of premature children with low-birth weight, a factor associated with increased risk of HB [4]. In Brazil, collected data are concordant with the HB world prevalence *(www.inca.gov.br/wcm/incidencia/2017)*. The overall 5-year survival rate of children with hepatoblastoma is 70% [5, 6]; however, patients who do not respond to standard treatment have very low survival rate [[7], [8], [9], [10]]. Few cases in adults have also been described [[11], [12], [13]], and prognosis is most unfavorable. The Children’s Hepatic Tumors International Collaboration (CHIC) has developed a novel risk stratification system on the basis of prognostic features present at diagnosis [14, 15]. Five backbone groups were defined according to clinical prognostic factors – age, AFP level (≤100 ng/mL), PRETEXT group (I, II, III, or IV), and metastasis at diagnosis.

HB genomes are relatively stable, with few cytogenetic alterations, mostly gains of chromosomes 2, 8 or 20 [[16], [17], [18]]. The major driver mutations in HB tumorigenesis are mainly activators of the WNT pathway, with recurrent mutations in *CTNNB1* [[19], [20], [21],[22]]. Few other molecular mechanisms engaged in HB tumorigenesis include overexpression of *IGF2* [23] and its transcriptional activator *PLAG1* [24], and down-regulation of *RASSF1A* by promoter hypermethylation [25]. This relative paucity of molecular biomarkers in HBs poses a challenge to proper stratification and adjustment of the therapeutic regimen, and molecular sub classification including gene signatures that could be used to stratify patients with hepatoblastoma were reported in the last years [[2], [19], [26]].

Exome sequencing has broadened the understanding of the HB mutational profile [[19], [27], [28], [29], [30]]. The commonalities disclosed by these studies, besides *CTNNB1* mutations, were the low number of detectable somatic mutations, and pathogenic variants in genes from the WNT pathway, such as *CAPRIN2* [27]. Other mutations were involved with the ubiquitin ligase complex *(SPOP, KLHL22, TRPC4AP*, and *RNF169)* [27], and with the transcription factor *NFE2L2*, impairing the activity of the KEAP1/CUL3 complex for proteasomal degradation [19, 28]. Clinically, overexpression of *NQO1*, a target gene of *NFE2L2*, was significantly associated with poor outcome, metastasis, vascular invasion, and the adverse prognostic C2 gene signature [26]. Other two exome analysis were based on syndromic patients who developed HB, including a boy with Simpson-Golabi–Behmel syndrome carrier of a germline *GPC3* loss of function (LoF) mutation (29), and a girl presenting severe macrocephaly, facial dysmorphisms and developmental delay, in which a novel *de novo* germline nonsense mutation was detected in the *WTX* [30]. In a recent study [31], 16 HBs were included in a pan-cancer cohort of pediatric tumors, with the identification of *CTNNB1* and *TERT*, genes already known to be frequently mutated in this type of tumor.

We describe here the mutational signatures and exome findings of a cohort of ten HBs, disclosing somatic mutations relevant as well as revealing a potential new biological mechanism, corroborated by expression and protein analyses. In addition, germline mutations were investigated in a rare patient with congenital HB.

## PATIENTS AND METHODS

### Patients

This study was approved by Research Ethics Committee – A.C. Camargo Cancer Center, (number 1987/14). Participants and / or persons responsible signed an informed consent form. All methods were performed in accordance with the relevant guidelines and regulations. Fresh-frozen tumor and matched non-tumoral liver tissues and blood samples were retrieved from ten hepatoblastoma patients of the A. C. Camargo Cancer Center Biobank (ten HB samples=exome cohort, five matched non-tumoral liver tissues, and five matched blood samples). A validation cohort was used for targeted sequencing, and RNA expression studies, comprising 12 additional HB cases (11 from GRAACC – Adolescent and Child with Cancer Support Group -, and one from A. C. Camargo Cancer Center; clinical features of this second HB cohort are described in **Supplementary Table 1**). All patients received pre-surgery chemotherapy according to both SIOPEL *(http://www.siopel.org/)* and COG *(https://www.childrensoncologygroup.org/)* protocols. This work was approved by the A. C. Camargo Cancer Center and GRAACC ethics committee; samples were collected after signed informed consent obtained from parents. Patients were followed by clinical examination, imaging tests and alpha-fetoprotein dosage.

In addition to the Brazilian HBs cohorts, a validation set of 16 additional HBs was tested (**Supplementary Table 1**; TCH samples). All these samples were de-identified specimens selected from the Texas Children’s Hospital Department of Pathology archives (Molecular Oncology Laboratory), after IRB approval (Baylor College of Medicine Institutional Review Board).

### DNA and RNA isolation

DNA and RNA were extracted from liver and blood samples following the technical and ethical procedures of the A.C. Camargo Tumor Bank [32, 33], using QIASymphony DNA Mini kit *(QIAGEN)* and RNeasy Mini Kit *(QIAGEN).* From tissue embedded in paraffin, direct cut (10 μg) and phenol-chloroform extraction were applied. Purity and integrity of DNA samples were checked by electrophoresis in 0.8% agarose gels and spectrophotometry (NanoDrop, *Thermo Scientific)*, and RNA samples were evaluated by microfluidics-based electrophoresis (Bioanalyzer, *Agilent Technologies);* only high-quality RNA samples (RIN >7.0) were used for gene expression analysis.

### Exome sequencing analysis

Exome data were obtained from genomic libraries of ten HBs and matched non-tumoral samples, enriched using the Sureselect 244K V3 *(Agilent Technologies*; n=11), OneSeq Constitutional Research Panel *(Agilent Technologies;* n=5), and QXT SureselectV6 *(Agilent Technologies;* n=4). Enriched libraries were sequenced on the Illumina HiSeq2500 platform using a 150-bp paired-end protocol to produce a median coverage depth on target of at least 50X per sample. Reads were mapped to their location in the human genome hg19/Grch37 build using the Burrows-Wheeler Aligner (BWA) package version 0.7.17. Local realignment of the mapped reads around potential insertion/deletion (indel) sites was carried out with the Genome Analysis Tool Kit (GATK) version 3.8. Duplicated reads were marked using Picard version 2.18, reducing false-positive SNP calls. Additional BAM file manipulations were performed with Samtools 1.7. Base quality (Phred scale) scores were recalibrated using GATK’s covariance recalibration. Somatic SNPs and indel variants were called using the GATK Mutect2 for each sample. A total of 53.43 Gigabases of sequence data were aligned at high quality (95% of aligned reads), with a mean of 4.45 Gb per sample. More than 95% of the sequenced bases presented Phred score >20. An average coverage depth of 42.6-fold per sample was achieved, with a median of 78% of target regions covered at a minimum of 20× read depth.

Data annotation and filtering variants were run through VarSeq software version 1.5.0 *(Golden Helix)* using the vcf. files. Variant annotation was performed using different public databases, including population frequency, such as EXAC *(http://exac.broadinstitute.org/)*, gnomAD (Genome Aggregation Database – *http://gnomad.broadinstitute.org/*), ABRaOM *(http://abraom.ib.usp.br/)*, 1000 genomes *(http://www.1000genomes.org/)*, and dbSNP version 138 *(http://www.ncbi.nlm.nih.gov/projects/SNP/)*; cancer mutation databases – COSMIC version 67 *(http://cancer.sanger.ac.uk/cancergenome/projects/cosmic/)* and ICGC *(http://icgc.org/)* -, and clinical sources – Clinvar (https://www.ncbi.nlm.nih.gov/clinvar) and OMIM *(https://www.omim.org)*. Variant filtering was based on quality (Phred score >17), read depth (>10 reads), variant allele frequency (>10%), population frequency (<0.001%), and predicted protein effect (missense, and LoF – essential splice site, frameshift and nonsense variants). *In silico* prediction of pathogenicity of missense variants were based on six algorithms provided by the database dbNSFP *(http://varianttools.sourceforge.net/Annotation/DbNSFP, version 2.4):* SIFT *(Sorting Intolerant from Tolerant- http://sift.bii.astar.edu.sg/*), Polyphen 2 *(Polymorphism Phenotyping v2 – http:/genetics.bwh.harvard.edu/pph2/)*, Mutation Taster *(http://mutationassessor.org/)*, Mutation Assessor *(http://mutationassessor.org/)*, and FATHMM *(Functional Analysis through Hidden Markov Models (V2.3) – http://fathmm.biocompute.org.uk/*). The potential damaging effect was also assessed using the VEP32script software package from Ensembl *(https://www.ensembl.org/)*. Likely pathogenic variants were visually validated as somatic alterations using both Integrated Genomics Viewer (IGV) and Genome Browser *(Golden Helix*).

Selected variants were validated by target sequencing; the gene panel was elaborated based on genes disclosed in the current exome analysis *(Agilent’s SureDesign* platform with a total of 18,539 probes and a total size of 498,019 kbp). Libraries were prepared from 22 fresh-frozen samples (exome and validation cohorts) using the *244K Agilent SureSelect Target Enrichment (Agilent Technologies)* system; the *TruSeq v2* chemistry 500 cycles kit was used with 250pb paired-end-protocol on the *Illumina MySeq*. The software *SureCall (Agilent Technologies)* was used for analysis.

### Sanger Sequencing

Prioritized variants from five candidate genes (from our study and the literature; *CTNNB1, CAPRIN2, CX3CL1, AXIN1*, and *DEPDC5)* were validated by Sanger sequencing (sequences upon request) in the HB exome cohort of ten tumors, and investigated in 24 additional samples (12 HBs of the validation group; and additional 12 HBs from formalin-fixed paraffin-embedded samples that were contained in a tissue microarray previously made in the Institution; the clinical information about the cases included in the tissue microarray is available in **Supplementary Table 2**). 14 HB cases from the Texas Children`s Hospital were screened for the *CX3CL1* variant. PCR reactions were performed using standard conditions (95 °C, 5 min; (44 °C, 30 s.; * °C, 30 s.; 72 °C, 45 s) × 30 cycle; 72 °C, 10 min), and amplicons were sequenced in both directions using an ABI 3730 DNA sequencer *(Applied Biosystems*); sequences were aligned with the respective gene reference sequence using Chromas Lite software *(Technelysium).*

### Gene expression analysis by real-time PCR

Gene expression was performed using the exome (n=9) and validation cohorts (n=10), and six liver tumor cell lines (HEPG2, C3A, SNU-387, SNU-423, SNU-449 and SNU – 475) for *CX3CL1* and *CX3R1.* RNA to cDNA conversion was made using the *Applied Biosystems* High Capacity RNA to cDNA kit following the manufacturer’s protocol. For qPCR, we used TaqMan Universal Master Mix II *(Applied Biosystems)* with reactions performed on an ABI PRISM 7500 instrument. *18S* was selected as the most stable reference gene among *18S, B2M, GAPDH* and *ACTA1* genes tested according to geNorm [34]. Averages from sample triplicates were compared between groups, considering differentially expressed those genes with fold changes ≥|2| through the 2^-ΔΔCt^ relative quantitative (RQ) method [35], with p-value ≤0.05. Mann-Whitney test was applied in the analysis of all tumors and cell lines compared to the control group; paired tumor/normal tissue samples were compared using the Wilcoxon test. All tests were corrected using Bonferroni. Prism 6 software *(GraphPad; CA, USA)* was used for statistical analyses.

### Immunohistochemistry

Protein analysis was performed for two genes *(CX3CL1*, and *CX3CR1)* using the following antibodies: *Polyclonal* Antibody PA5-23062 *(CX3CL1)*, and Polyclonal Antibody PA5-19910 *(CX3CR1)*, both from *ThermoFisher* scientific company. Reactions were automated in the BenchMark Ultra-VENTANA equipment or manual protocol *(Novocastra Novolink kit).* In total, immunohistochemistry were evaluated in 34 cases: nine HB samples from the exome cohort, 17 additional HBs from the tissue microarray [36] and eight samples from the Texas Children’s Hospital cohort, including a lung metastasis sample.

### Mutational signature detection

Exome data of HBs and matched non-tumoral tissues were used to detect specific mutational signatures. All somatic single base substitutions were mapped onto trinucleotide sequences by including the 5` and 3’ neighboring base contexts to construct a 96 x G matrix of mutations count. Next, we used signeR [37] to estimate the number of mutational processes and their signatures. Cosine similarity score was used to compare the signatures with the Pan-Cancer catalog of 30 signatures in COSMIC database.

## RESULTS

Clinical features of the cohort of the ten HBs that were studied by exome sequencing are described in **Table 1**. The mean age at diagnosis was 24.5 months, excluding one patient who was diagnosed at 17 years (HB28). One of the patients was syndromic (born premature, underweight, developmental delay, facial dysmorphisms, and craniosynostosis; HB46), and two others presented kidney abnormalities, including the patient that developed a congenital tumor (HB33). Four cases were classified as high-risk according to CHIC criteria, and three of these patients presented pulmonary metastasis at diagnosis. After histopathological reexamination, one case was reclassified as presenting HB/Hepatocellular Carcinoma (HCC) features (HB30). Three patients died from the disease, including the patient diagnosed at 17 years old and the case reclassified as HB/HCC features; the third patient (HB15) died from complications of liver transplantation.

**Table 1.** Clinical features of ten hepatoblastoma cases investigated by exome sequencing.

| ID/gender/ age at diagnosis | Histology | AFP ng/ml | Risk stratification*/PRETEXT | Chemotherapy Protocol | Transplant | Metastasis | Relapse | Overall Survival | Premature (low birth weight) | Other features | Type of analysis |
| --- | --- | --- | --- | --- | --- | --- | --- | --- | --- | --- | --- |
| HB15, F, 18m | Epithelial Embryonal | 5668000 | Intermediate/4 | NA | Yes | No | No | 1 year | No | . | Exome sequencing, mutation screening by Sanger sequencing, RNA expression and IHC assays |
| HB16, M, 9m | Epithelial Fetal | 824 | Intermediate/4 | SIOPEL3 | No | No | No | >5 years | No | . | Exome sequencing, mutation screening by Sanger sequencing and IHC assays |
| HB17, F, 36m | Epithelial Fetal | >400000 | Low/1 | SIOPEL3 | No | No | No | >5 years | No | . | Exome sequencing, mutation screening by Sanger sequencing, RNA expression and IHC assays |
| HB18, M, 9m | Epithelial and Mesenchymal mixed | >200000 | Low/3 | SIOPEL3 | Yes | No | No | >5 years | No | . | Exome sequencing, mutation screening by Sanger sequencing, RNA expression and IHC assays |
| HB28, M, 17y | Epithelial and Mesenchymal mixed | NA | High/4 | SIOPEL4 | No | No | Yes | 4 years | No | Hepatomegaly at birth | Exome sequencing, mutation screening by Sanger sequencing and RNA expression |
| HB30, M, 54m | HB with HCC features - Epithelial Fetal | >1000000 | High/2 | SIOPEL4 | Yes | Lung | Yes | 5 years | No | . | Exome sequencing, mutation screening by Sanger sequencing, RNA expression and IHC assays |
| HB31, M, 30m | Epithelial Fetal | 742000 | Low/3 | NA | No | No | No | >5 years | No | Non-functional kidney | Exome sequencing, mutation screening by Sanger sequencing, RNA expression and IHC assays |
| HB32, F, 36m | Epithelial and Mesenchymal mixed | 9328000 | High/4 | SIOPEL4 | Yes | Lung | No | >5 years | No | . | Exome sequencing, mutation screening by Sanger sequencing, RNA expression and IHC assays |
| HB33, F, 1m | Epithelial Embryonal and Fetal | 28312000 | Intermediate/2 | SIOPEL3 | No | No | No | >5 years | No | Congenital HB and renal agenesis | Exome sequencing, mutation screening by Sanger sequencing, RNA expression and IHC assays |
| HB46, M, 28m | Epithelial and Mesenchymal mixed | >200000 | High/4 | SIOPEL6 | No | Lung | No | >3 years | Yes | Syndromic patient # | Exome sequencing, mutation screening by Sanger sequencing, RNA expression and IHC assays |

Caption: F: female; M: male; NA: data not available; AFP: Alphafeto protein; IHC: Immunohistochemistry
\*According to the CHIC criteria (15)

### Identification of somatic coding non-synonymous mutations by exome sequencing

The strategy of analysis of the exome sequencing data was designed to identify somatic variants, excluding non-coding and coding synonymous variants. Only LoF and missense somatic mutations, the later predicted to be pathogenic by at least one *in silico* algorithm, were considered in this analysis. A total of 94 somatic non-synonymous mutations were disclosed (92 different variants), mapped to 87 different genes (**Supplementary Table 3**); the detected mutations were validated either by targeted or Sanger sequencing. Two HBs did not present detectable somatic non-synonymous coding mutations (HB17 and HB28), and another one (the congenital case HB33) was found to harbor 40% of the identified somatic mutation in this cohort. The mean number of somatic non-synonymous mutations per sample was 9.4; however, excluding the atypical sample (HB33), the median number of somatic nonsynonymous variants was 6.2 per tumor.

Four of these detected mutations were already reported in COSMIC, three of them in *CTNNB1* (c.101G> A: COSM5671; c.101G> T: COSM5670; c.121A> G: COSM5664), and one in *GMPS* (c.1367G>T: COSM1040323). The same missense mutation (c.704C>G, A235G) was disclosed in the *CX3CL1* gene in two tumors (HB32 and HB33). *CEP164* gene was mutated in two tumors, although presenting different variants. Table 2 presents details of the mutations considered to be likely pathogenic: six LoF variants (five nonsense and one frameshift, five of them in a single tumor), and six missense variants (recurrent variants or recurrent genes in different tumors). A validation cohort of 12 HBs was screened for the full set of somatic variants, and only *CTNNB1* mutations were detected.

**Table 2:** Description of loss-of function, recurrent genes and variants somatically identified in ten hepatoblastomas by exome sequencing (genomic coordinates according to the GRCh37/hg19 Human Assembly): variant data#, mutation type, effect on protein, and prediction of pathogenicity.

| ID | Gene | Chr:genomic coordinate (rs) | VF (%) | RefSeq | Variant type | AA Change | Protein change | Pathogenicity score* |
| --- | --- | --- | --- | --- | --- | --- | --- | --- |
| HB15 | *CEP164* | 11:117258055 | 14 | NM_014956 | missense | c.1861C>A | p.Leu621Met | 2/5 |
| HB31 | *CEP164* | 11:117267312 | 17 | NM_014956 | missense | c.3263A>G | p.Asp1088Gly | 2/5 |
| HB16 | *CTNNB1* | 3:41266104 | 21 | NM_001098210 | missense | c.101G>A | p.Gly34Glu | 3/5 |
| HB33 | *CTNNB1* | 3:41266104 (rs28931589) | 58 | NM_001098210 | missense | c.101G>T | p.Gly34Val | 3/5 |
| HB46 | *CTNNB1* | 3:41266104 (rs28931589) | 52 | NM_001904 | missense | c.101G>A | p.Gly34Glu | 4/5 |
| HB18 | *CTNNB1* | 3:41266124 (rs121913412) | 43 | NM_001904 | missense | c.121A>G | p.Thr41Ala | 3/5 |
| HB32 | *CX3CL1* | 16:57416454 | 11 | NM_002996 | missense | c.704C>G | p.Ala235Gly | 2/5 |
| HB33 | *CX3CL1* | 16:57416454 | 40 | NM_002996 | missense | c.704C>G | p.Ala235Gly | 2/5 |
| HB31 | *ACACA* | 17:35581924 | 24 | NM_198834 | stop codon | c.3463G>T | p.Glu1155Ter | . |
| HB33 | *ARVCF* | 22:19960467 | 35 | NM_001670 | stop codon | c.2531C>T | p.Trp844* | 1/5 |
| HB33 | *DEPDC5* | 22:32215040 | 40 | NM_001242896 | stop codon | c.1699C>T | p.Arg567* | 1/5 |
| HB33 | *MYH7* | 14:23893250 | 17 | NM_000257 | stop codon | c.2788G>T | p.Glu930Ter | 5/5 |
| HB33 | *NOL6* | 9:33466636 | 17 | NM_022917 | stop codon | c.2022C>T | p.Trp674* | 3/5 |
| HB33 | *KIAA0319L* | 1:35900602 | 29 | NM_024874 | frameshift | c.3042*>+T | p.Phe1014X | 1/5 |

Caption: ID: Identification of the sample in the project; VF: Frequency of the variant allele; RD: Read Depth; AA: aminoacid
\*The pathogenicity score indicates the number of algorithms that predicted for a given missense variant to be deleterious to the protein function (Polyphen2, SIFT, Mutation Taster, Mutation Assessor Pred, FATHMM Pred) .

Additionally, an integrative analysis based on previous DNA methylation (DNAm) data from the same group of HB samples [38] showed a partial overlap between the set of genes presenting somatic mutations and the set with DNAm changes: *EGFR* and *LMBRD1*, hypermethylated, and *AHRR*, hypomethylated, respectively.

To explore the pathways in which the mutated genes are involved and their biological roles, we used KEGG *(Kyoto Encyclopedia of Genes and Genomes http://www.genome.jp/kegg*; Release 86.1, May 10, 2018) and Gene Ontology *(http://www.geneontology.org/*; PANTHER Over representation Test, Homo sapiens – REFLIST 21042) databases. It was detected an enrichment for several development processes, Pathways in cancer, Proteoglycans in cancer, Metabolic pathways, Cytokine-cytokine receptor interaction, among others.

### Known and novel CTNNB1 mutations

We investigated the presence of *CTNNB1* mutations either by exome or Sanger sequencing in the ten HBs and additional 12 tumors from the validation group. A total of seven pathogenic variants were detected in eight samples. Six *CTNNB1* missense mutations (G34E, G34V, T41A, D32A, S29F, and S33C; **Figure 1a-e**), which had already been reported in HBs (COSMIC), and a novel likely pathogenic variant, a 39 bp inframe deletion (A21_S33del) (**Figure 1f** – HB40T). All identified variants map in the *CTNNB1* exon 3 (**Figure 1e**), at GSK3β phosphorylation sites. Additionally, six tumors presented size-variable *CTNNB1* intragenic deletions that were ascertained by Sanger sequencing. In summary, *CTNNB1* alterations were detected in 14 out of the 22 tested HBs (64%).

**Fig 1:**
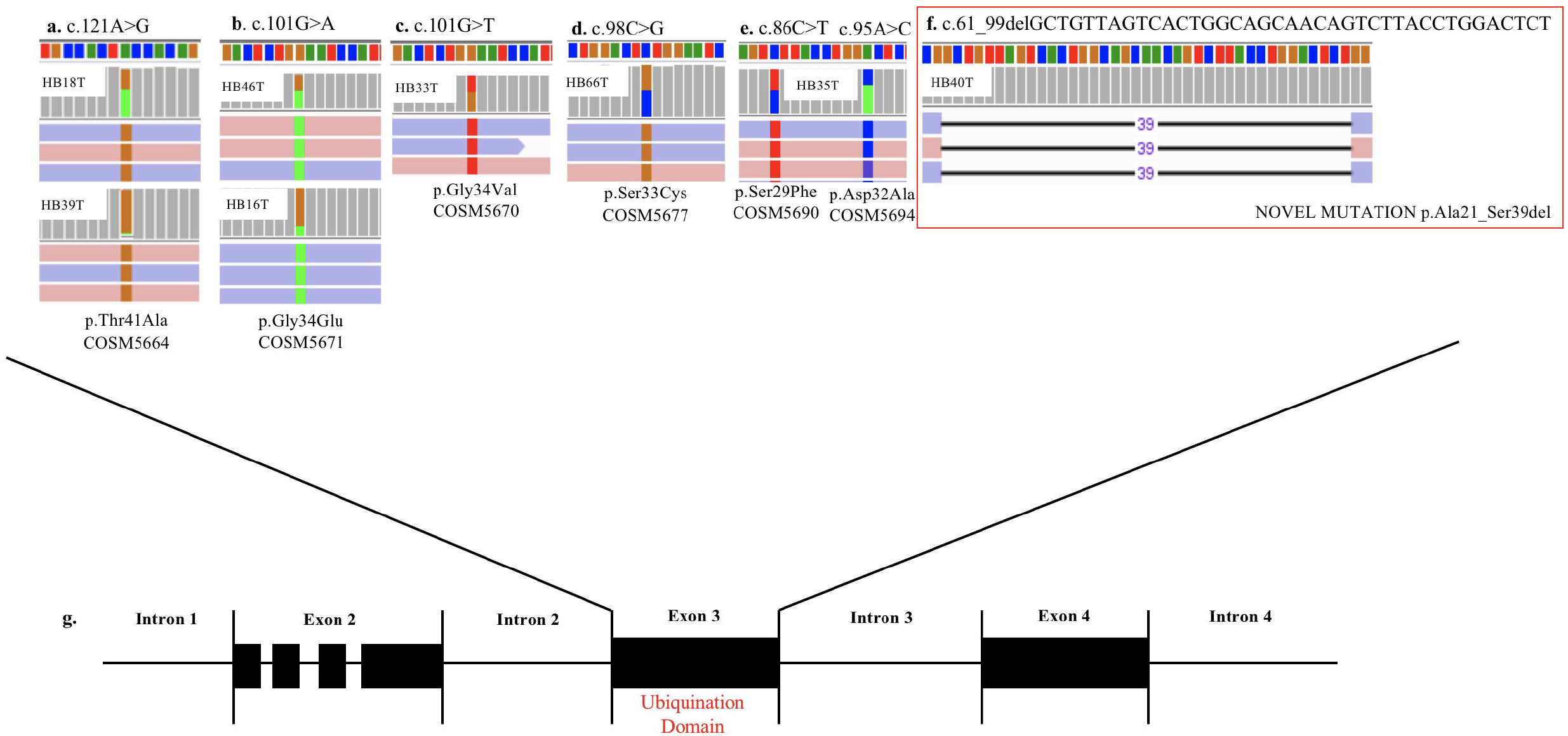
*CTNNB1* somatic mutations detected in eight hepatoblastoma samples: The upper panel presents the six different *CTNNB1* somatic mutations identified by exome sequencing in eight tumors; BAM file images from tumor NGS data show mutations which were detected in both directions (pink and blue bars correspond to forward and reverse reads, respectively): **A**. HB18T (variant frequency of 43%) and HB39T (variant frequency of 11%), mutation c.121A> G; **B**. HB46T (variant frequency of 52%) and HB16T (variant frequency of 21%), mutation c.101G>A. **C**. HB33T (variant frequency of 58%), mutation c. 101G>T. **D**. HB46T (variant frequency of 50%), mutation c.98C>G. **E**. HB35T, two mutations: c.86C>T (variant frequency of 49%) and c.95 A>C (variant frequency of 44%); **F**. HB40T, the novel *CTNNB1* likely pathogenic variant reported in the present study: a 39bp inframe deletion c.61_99delGCTGTTAGTCACTGGCAGCAACAGTCTTACCTGGACTCT (variant frequency of 21%). **G**. Detected mutations are all mapped in the exon 3 of the gene, at the ubiquination domain.

### Recurrent A235G somatic mutation in CX3CL1: a new HB gene?

The missense mutation C>G at the position 704 of the exon 3 of *CX3CL1* (NM_002996) was identified in two samples (HB32 and HB33). This mutation leads to substitution of the amino acid alanine by glycine in the codon 235 of the protein, predicted as damaging for protein function by SIFT and Mutation Taster algorithms (**Figure 2a**). The *CX3CL1* variant was validated by target sequencing in both mutated tumors (**Figure 2b**); Sanger sequencing detected the mutation only in the tumor sample with the higher variant frequency (HB33, 40%). Another 47 HB samples were also tested for the presence of the *A235G* variant, but no mutations were identified.

**Fig 2.**
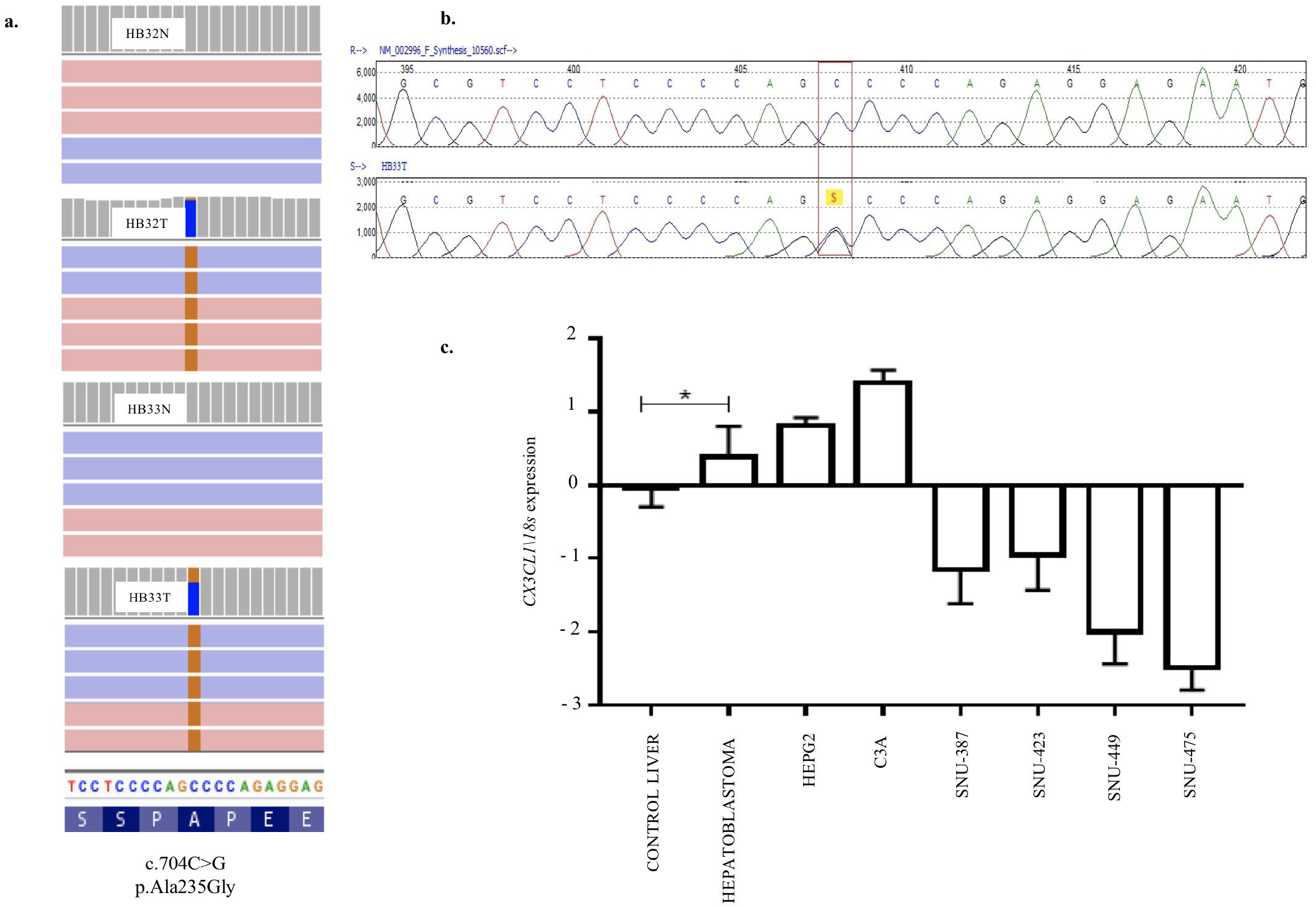
A recurrent A235G somatic mutation detected in the exon 3 of the *CX3CL1* gene and pattern of RNA expression in hepatoblastomas: **A**. Image obtained from IGV; BAM file images from tumors (HB32T and HB33T) and germinative non-tumoral (HB32N and HB33N) samples showing that the A235G mutations (c.704C>G, p.Ala235Gly) were detected in both directions (pink and blue bars correspond to forward and reverse reads, respectively); HB32T exhibiting a low variant frequency (11%) and HB33T with a variant frequency of 40%. **B**. Sanger Sequencing showing the *CX3CL1* variant in heterozygosity. c. Gene expression pattern of the *CX3CL1* gene in 19 HB samples; HB samples, including the CX3CL1-mutated HB32 and HB33 tumors, and the HB cell lines (HEPG2 and C3A) presented upregulation in comparison to control liver samples. The hepatocellular carcinoma cell lines (SNU-387, SNU-423, SNU-449 and SNU-475) were found to be down-regulated in relation to control samples and HBs. The statistical test used was Mann-Whitney, *p<0.05 (Bonferroni correction); Endogenous gene: *18s* and the controls are non-tumoral liver tissues. For the analyzes the values in log of RQ were used.

*CX3CL1* expression level was evaluated in 19 HB samples (including the two mutated ones), nine non-tumoral liver samples, two hepatoblastoma cell lines, and four hepatocellular carcinoma cell lines. Up-regulation of *CX3CL1* was detected in the HB group, including *CX3CL1-*-mutated tumors and HB cell lines, compared to control liver samples (fold-change >2; p<0.05) (**Figure 2c**). The hepatocellular carcinoma cell lines were found to be down-regulated in relation to control samples and HBs. To investigate if the presence of the *CX3CL1* mutation and/or up-regulation of its mRNA could influence the involved pathway, the expression of the *CX3CL1* receptor *(CX3CR1)* was also assessed. Six tumors presented upregulation of *CX3CR1* mRNA in relation to control (fold-change > 2), including one of the *CX3CL1-*-mutated tumors (HB32; **Supplementary Figure 1**). Vascular invasion was the only clinical characteristic with a trend towards *CX3CL1* upregulation (64% of high-expression samples *versus* 25% of low-expression; p<0.06, Chi-square test; data not shown). Expression of *CX3CR1* and *CX3CL1* did not seem to be correlated.

DNA methylation (DNAm) data at gene bodies and promoters of *CX3CL1* and *CX3CR1* were recovered for HB samples [38] to correlate with the expression level of these genes (**Supplementary Figure 2**). A significant DNAm decrease was observed in *CX3CL1* promoter in tumors compared to control liver samples *(p-adj* 0.006), and an inverse correlation between gene expression and DNAm level in the *CX3CL1* gene body (Spearman’s rho 0.46, P-value 0.02), although the latter presented great inter-tumor heterogeneity.

Protein analysis showed positivity of CX3CL1 in most of the investigated samples (20 out 26 HBs), presenting nuclear or cytoplasmatic labeling at different degrees (details available in **Supplementary Table 4**). HB32-mutated tumor exhibited weak cytoplasmatic labeling and nuclear positivity in more than 50% of cells, while HB33-mutated showed strong cytoplasmatic labeling and nuclear positivity in 25% of cells (**Figure 3a1-c1**); in particular, HB33 exhibited great heterogeneity of histology and protein labelling. Positive labeling of CX3CR1 was detected only in the two CX3CL1-mutated tumors (**Figure 3a-c**); HB33 showed cytoplasmatic signal, and HB32 had both nuclear and cytoplasmatic labeling. Non-tumoral liver samples did not show any labeling.

**Fig 3:**
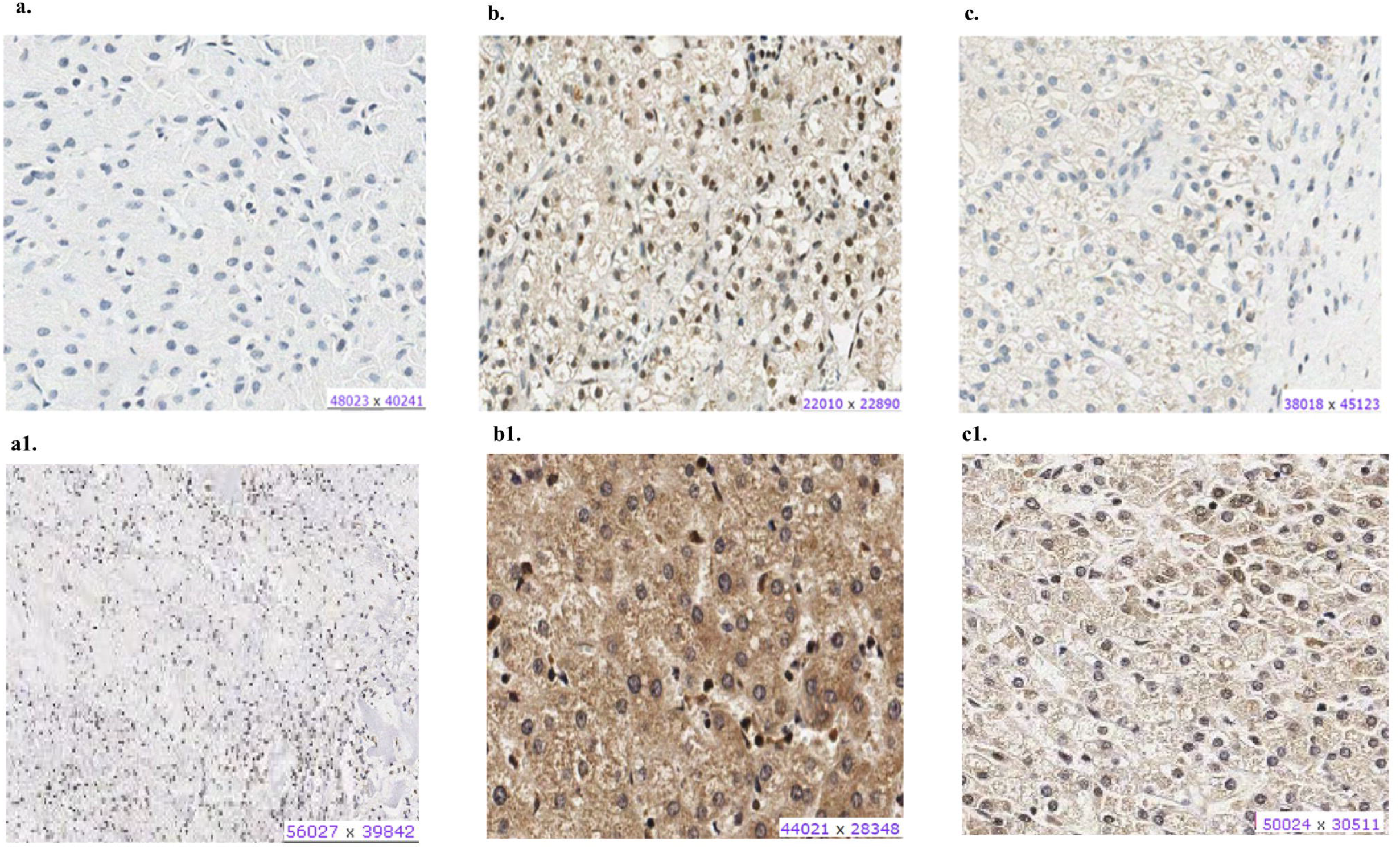
Protein expression of CX3CL1 and CX3CR1 evaluated in hepatoblastoma samples by immunohistochemistry assay: Panels **A-C** shows CX3CR1 data, and panels A1-C1, CX3CL1 from the same tumor samples. **A**. HB17, example of negative labeling for CX3CR1 **(A)** and CX3CL1 **(A1)**. **B**. HB32T, positive for nuclear and cytoplasmic CX3CR1 **(B)** and CX3CL1 **(B1)**. **c**. HB33T, positive for cytoplasmic CX3CR1 labeling **(C)** and positive for nuclear and cytoplasmic CX3CL1 **(C1)**.

An independent set of eight HBs and one HB lung metastasis was also evaluated by immunohistochemistry, in a qualitative analysis; similarly to our previous observation, the pattern of protein expression indicated an activation of the CX3CL1/CX3CR1 pathway, with a predominance of these proteins in the cytoplasm of tumor cells (**Supplementary Table 5**). It was observed that in inflammatory regions of the tumors, both proteins were not expressed in the infiltrated lymphocytes, in which they should be expressed in normal conditions, whereas in necrotic regions, the protein staining was negative in tumor cells, but strongly positive in the infiltrated lymphocytes. Figure 4 illustrates these findings.

**Fig 4:**
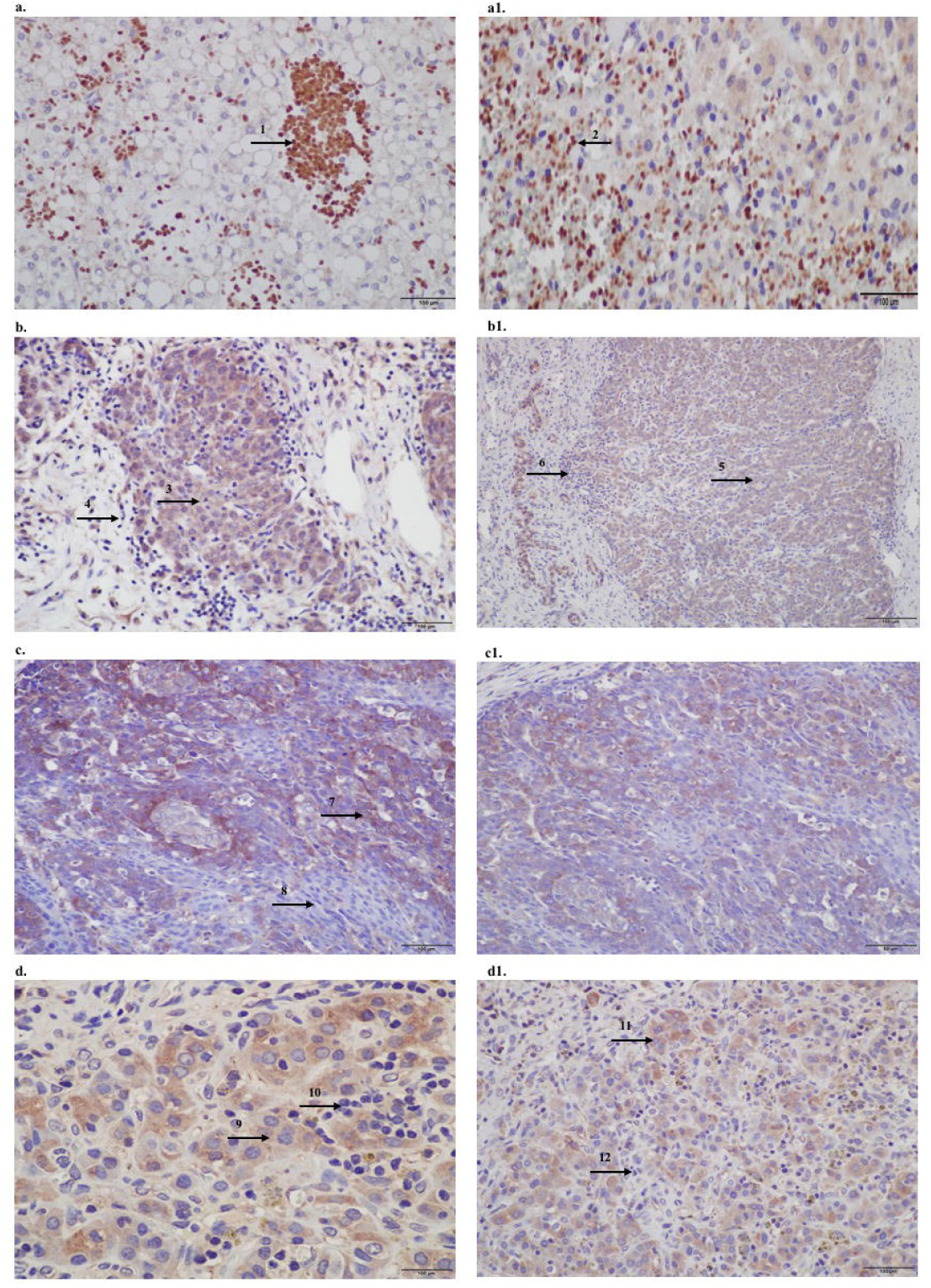
Protein expression of CX3CL1 and CX3CR1 evaluated in hepatoblastomas and hepatoblastoma lung metastasis by immunohistochemistry assay: Panels **a-d** shows CX3CL1 data, and panels a1-d1, CX3CR1. **A**. TCH361, CX3CL1 strongly positivity of infiltrated lymphocytes (indicated by arrow 1) in necrotic regions of the tumor, and **A1**. CX3CR1 strongly positivity of infiltrated lymphocytes (indicated by arrow 2) in necrotic regions of the tumor; **B**. and **B1**. TCH327, positivity in tumor cells (indicated by arrows 3 and 5) and infiltrated lymphocytes negative (indicated by arrows 4 and 6) for both proteins. **C**. TCH321, positivity in the osteoblast component and strong positivity in the fetal type (indicated by arrow 7); infiltrated lymphocytes are negative (indicated by arrow 8); C1. Positivity in tumor cells and lymphocytes negative; **D**. and **D1**. TCH360, lung metastasis showing positivity in tumor cells (indicated by arrows 9 and 11), and no expression in infiltrated lymphocytes (indicated by arrows 10 e 12), for both proteins.

### Mutational signatures of HB

Three signatures were detected (HB-S1, S2, and S3), two of them (HB-S1 and HB-S2) presenting great superposition to mutational signatures already reported in COSMIC. The profile of each signature is displayed in **Figure 5** using the six substitution subtypes (C>A, C>G, C>T, T>A, T>C, and T>G). HB-S1 group was most similar to COSMIC signatures 1 and 6, and HB-S2 group presented features of the COSMIC signature 29. HB-S3 showed no clear similarity to any known signature, presenting an unspecific pattern with a slight increase of C>A mutations. The relative signature contribution to mutations in each hepatoblastoma sample can be found in **Supplementary Figure 3**.

**Fig 5:**
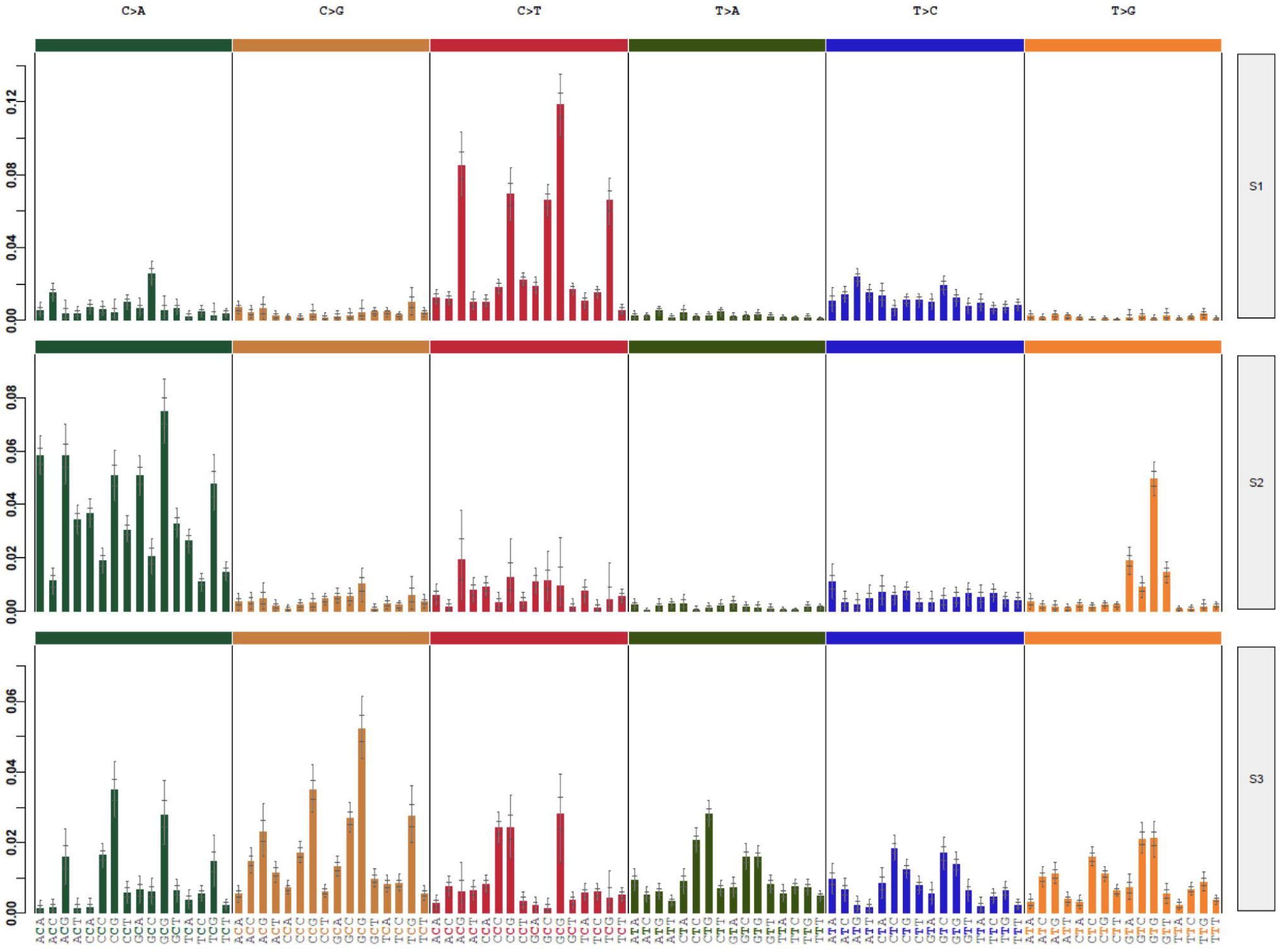
Three different mutational signatures were identified in hepatoblastomas: Exome data of HBs and matched germline tissues were used to detected specific mutational signatures (37). The profile of each signature is displayed using the six substitution subtypes (C>A, C>G, C>T, T>A, T>C, and T>G).

### Congenital HB case

Germline exome analysis was performed for this patient and her mother; father was unavailable. A total of 144 rare germline non-synonymous variants were identified in the patient, and absent from her mother (information on frequency and pathogenicity scores of the detected variants are available in **Supplementary Table 6**). Twelve germline variants were LoF *(AARSD1, ACSM3, ERI2, CECR2, CRYGA, DNAH7, ETV4, HOXC4, MAMDC4, NEBL, PRSS56*, and *TBXAS1)*, standing out a stop gain in *HOXC4*, which was not previously reported in any germline database, including a cohort of Brazilians (ABRAOM), and an indel in the *PRSS56* gene (ClinVar 31077), both variants already reported in liver cancer samples (ICGC). Additionally, the patient carries six missense variants which were predicted to be deleterious for protein function using six different algorithms; among them, a variant affecting *BRCA1*, and two others not documented in any database *(GOLGA5* and *FAH* gene, **Supplementary Figure 4**).

## DISCUSSION

Our exome findings revealed a low mutational background in HBs, corroborating previous works [[19], [26], [31]], with only three genes presenting recurrent mutations, namely *CTNNB1, CX3CL1*, and *CEP164. CTNNB1* somatic mutations were detected in ~60% of the tumors here studied, including a novel pathogenic variant (A21_S33del). Mutation in the *A2ML1* gene also appeared in common between our HB cohort and one of the major exome studies of HB samples [28], although the role of *A2ML1* somatic mutations remains unclear. Mutations in the promoter of the *TERT* gene was also reported as recurrently mutated in HBs; however, *TERT* promoter was not covered in this exome data.

Our data pointed out to a novel set of candidate genes for HB tumorigenesis. *CEP164*, a key element in the DNA damage-activated signaling cascade [39] involved in genome stability, was found to be mutated in two different HBs. *CEP164* is overexpressed in various cancer types, often associated with poor prognosis [40], and a recent study in rhabdomyosarcoma cells suggested a central role of this gene in proliferation in response to cellular stress [41]. Remarkably, one of the CEP164-mutated HBs exhibited a complex genome, with several copy number alterations and two large LOH regions. Three genes, which we have previously reported as differentially methylated in HBs [38], were mutated in the present cohort, reinforcing a possible role in HB tumorigenesis: *EGFR, LMBRD1*, and *AHRR. LMBRD1* encodes a lysosomal membrane protein and is associated with a vitamin B12 metabolism disorder [42], and *AHRR* and *EGFR* are involved in regulation of cell growth and differentiation. Six LoF variants were identified in *ACACA, ARVCF, DEPDC5, MYH7, NOL6* and KIAA0319L. Nevertheless, all but the *ACACA* variant were detected in the congenital tumor, making difficult to associate these mutations with HB in general.

The most significant finding in this study was the detection of a recurrent somatic missense mutation at the *CX3CL1* gene. This gene, chemokine ligand 1 (C-X3-C motif), encodes a large transmembrane 373aa multiple-domain protein from the chemokine family, the fractalkine. This protein is present in endothelial cells of diverse tissues, such as brain, kidneys, and liver [43], and is related to leukocytes movement, including migration to inflammation sites [44, 45]. The cell adhesion and migration functions are promoted through interaction of fractalkine with the chemokine receptor CX3CR1, a transmembrane protein known to provide pro-survival signaling for anti-inflammatory monocytes, but also present in NK cells and T cells [46]. The mutation is located in a region that exerts a key role related to the binding to CX3CR1. Under inflammatory response conditions, cleavage of CX3CL1 by metalloproteinases generates a soluble chemokine, which binds to CX3CR1 in nearby cells and can induce adhesion, cell survival, and migration. The significance of *CX3CL1* mutations in cancer is yet poorly understood, but mutations in different types of tumor are reported, predominantly in gastric cancer (COSMIC) and hepatocellular carcinomas (TCGA). Gastric tumors exhibit increased *CX3CL1* expression [47], and the CX3CR1 receptor is highly expressed in association with more advanced stages. Xu et al. [48] and Yang et al. [49] published results of another chemokine *(CXCL5)* in liver cancer, with data also indicating an oncogenic role.

Expression studies showed *CX3CL1* upregulation in hepatoblastomas, a result that was corroborated by immunohistochemistry assays. We also observed increased *CX3CL1* expression in several HBs without detectable *CX3CL1* mutations, suggesting alternative pathways for its activation, such as the significant hypomethylation at the *CX3CL1* promoter disclosed in HBs. *CX3CR1* expression was increased in only part of the tumors, but it was noteworthy that only the two *CX3CL1-*-mutated tumors presented CX3CR1 protein, evidencing an activation of this chemokine pathway. Our results indicate that the activation of the CX3CL1-CX3CR1 pathway could be related to HB progression. Inappropriate expression or regulation of chemokines and their receptors are linked to many diseases, especially those characterized by an excessive cellular infiltrate, such as rheumatoid arthritis and other inflammatory disorders. In recent years, the involvement of chemokines and their receptors in cancer, particularly metastases, has been well-established [50, 51]. Chemokines produced serve to recruit leukocytes, which produce other cytokines, growth factors, and metalloproteinases that increase proliferation and angiogenesis. The metastasis process is facilitated by the regulation of particular chemokine receptors in tumor cells, which allows them to migrate to secondary tissues where the ligands are expressed [52]. In an independent HB group, a contrasting pattern of CX3CL1 and CX3CR1 was observed in regions of inflammation in the samples, and in areas with necrosis. Around necrotic regions, CX3CL1 and CX3CR1 were detected in the infiltrated lymphocytes, indicating a normal immune response; however, in inflammation regions both proteins were strongly positive in tumor cells and not detected in infiltrated lymphocytes, suggesting a mechanism of regulation of this pathway in favor of HB cells. This result further adds to previous studies showing that the activation of the ligand and receptor in chemokines may be involved in tumor invasion [[47], [48], [49]]. All these pieces of evidences reinforce the importance of the study of chemokines in tumors in general, and in HBs in particular.

In addition to revealing coding somatic mutations in HBs, exome data was used to search for mutational processes. In general, it was remarkable that the mutational signatures already reported specifically for liver cancer were not observed in these HBs, suggesting distinct mutational mechanisms for hepatocellular carcinomas and liver embryonal tumors. Two of the three different mutational signature here observed have superposition mainly with three known signatures from COSMIC (signatures 1, 6 and 29). Signature 29 has been observed only in gingiva-buccal oral squamous cell carcinoma developed in individuals with a tobacco chewing habit; this signature indicates guanine damage that is most likely repaired by transcription-coupled nucleotide excision repair. Among the several chemicals in smokeless tobacco that have found to cause cancer [55], the most harmful carcinogen are nitrosamines, which level is directly related to the risk of cancer and that can be also find in food such as cured meat, smoked fish and beer. Interestingly, O(6)-methylguanine detected in human cord blood in mothers highly exposed to such products implicates Nitrosodimethylamine exposure of the fetus and toxicity from dietary sources of these compounds [56]. Maternal dietary exposure to N-nitroso compounds or to their precursors during pregnancy has also been associated with preterm birth [57] and risk of childhood cancer [58]. Childhood cancer is most probably the combinatorial result of both genetic and environmental factors, and these networks between fetal exposure to environmental carcinogens such as nitrosamines from tobacco and/or dietary sources, preterm birth, and increased risk of childhood cancer may be an underlying cause for at least a subset of HBs. Finally, a subset of tumors, including two patients who died from the disease, exhibited a mutational pattern with no clear similarity to any known signature.

As a final point, we analyzed the germline exome of the patient with a congenital HB and renal agenesis, who developed a tumor exhibiting a heterogeneous histology. This tumor presented the highest number of somatic mutations herein detected, including *CX3CL1* and *CTNNB1* mutations, and its chromosome copy number profile was complex compared to the HB group (data not shown). In addition to very rare germline variants in genes related to liver function, such as *HOXC4, PRSS56* and *CYP1A1*, the patient carried two variants strongly predicted to be deleterious affecting *BRCA1* and *FAH*, both genes associated with cancer predisposition [59]. In particular, the *FAH* gene encodes a fumarylacetoacetate hydrolase enzyme that is mainly abundant in liver and kidneys [60], and germline mutations were already reported to increase the risk of hepatocellular carcinoma [61], although only in a recessive mode of inheritance.

Several lines of evidence indicate that childhood and adult cancers are distinct entities. In spite of intensive efforts, relevant genetic factors remain difficult to be captured in rare cancers such as embryonal tumors like HB. In summary, in this study, we provide new evidences that the activation of the CX3CL1/CX3CR1 chemokine signaling pathway can be involved in hepatoblastoma tumorigenesis or progression. We present the first assessment of mutation signatures in hepatoblastomas identifying a novel signature specific to a subset of these tumors. Additionally, we draw attention to the aspect of a likely strong genetic component of cancer predisposition at least in part of the HB patients, possibly related to the presence of additional clinical signs such as kidney abnormalities.

## ACKNOWLEDGEMENTS

We thank patients their families for participating in the study.

## COFLICTS OF INTEREST

We declare that we have no conflicts of interest.

## AUTHOR CONTRIBUTIONS

Conception and design: TFMA, TR, CR, ACVK

Collection and assembly of data: TFMA, MPR, SC, TR, JSB, ACB, AM, SFS, MC, SRCT, MLPA, DMC, CR, CMLC, IWC, DLT, ACVK

Data analysis and interpretation: TFMA, MPR, SC, TR, JSB, ACB, MM, RV, GRF, SFS, DLT, IT, DMC, CR, CMLC, IWC, ACVK

Manuscript writing: TFMA, CR, ACVK

Final approval of manuscript: All authors

