## Supplemental material for "Mutational burden of hepatoblastomas: a role for the *CX3CL1/CX3CR1* chemokine signaling pathway"

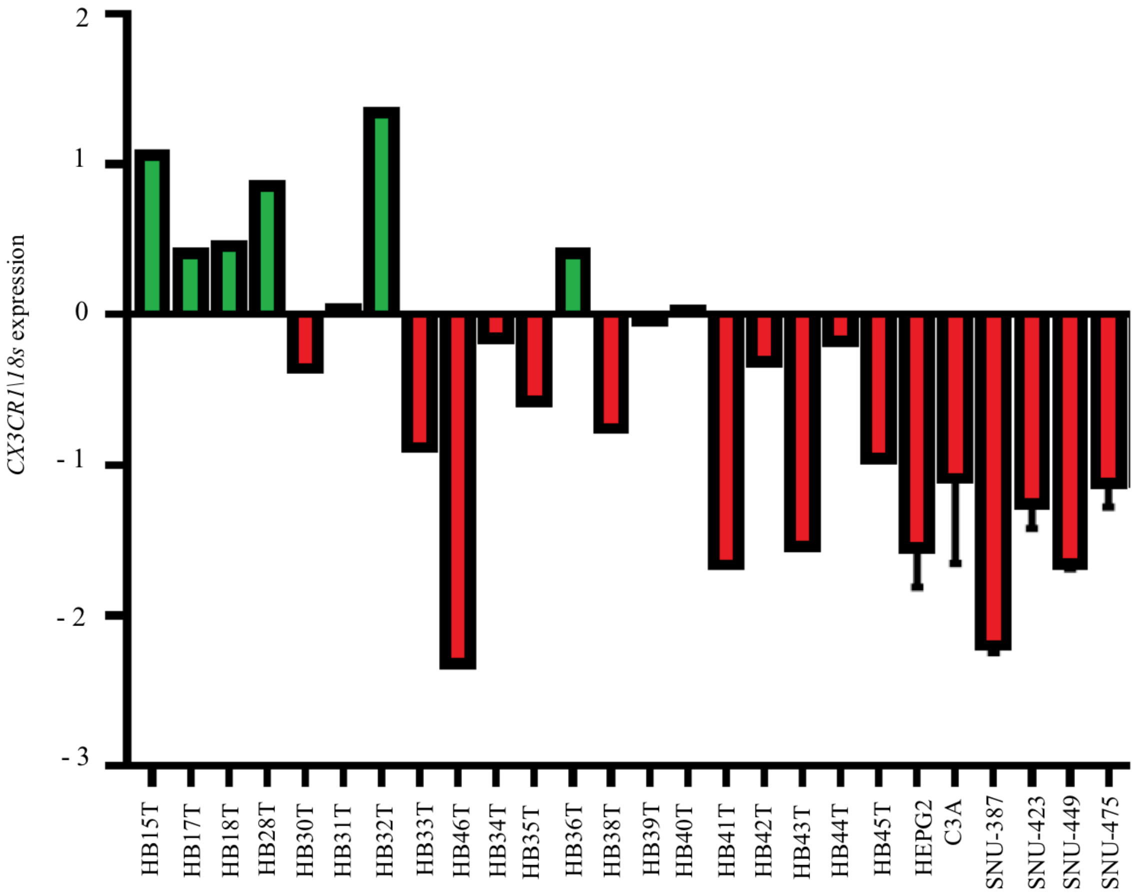


**Supplementary Figure 1: Gene expression pattern of the *CX3CR1* gene in 19 HB samples and six liver tumor cell lines.** Only six tumors (green bars) presented *CX3CR1* upregulation in comparison to control liver samples. The hepatoblastomas cell lines (HEPG2 and C3A) and the hepatocellular cell lines (SNU-387, SNU-423, SNU-449 and SNU-475) were found to be down-regulated in relation to control samples. The statistical test used was Mann-Whitney, no significance; Endogenous gene: 18s and the controls are non-tumoral liver tissues. For the analyzes the values in log of RQ were used.

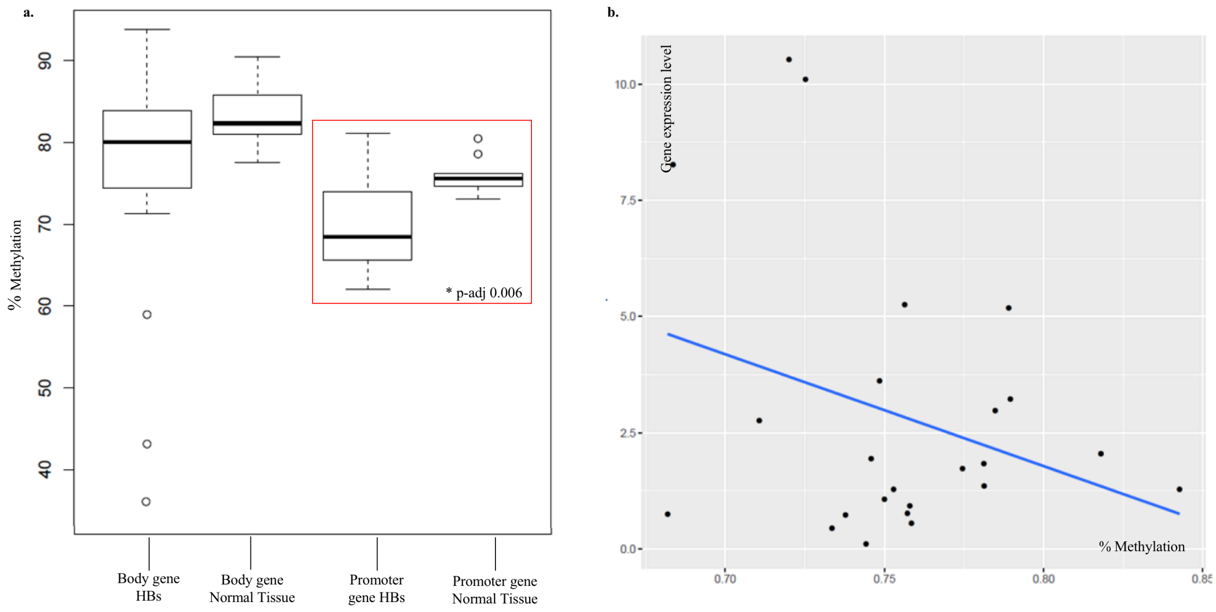


**Supplementary Figure 2: *CX3CL1* DNA methylation data recovered for the same set of HB samples from our previous work (38) and correlation with gene expression data: a.** A significant DNAm difference was observed in *CX3CL1* gene promoters of HBs in relation to control livers (p-adj 0.006). **b.** An inverse correlation between gene expression and DNAm level was detected in *CX3CL1* gene body (Spearmans rho 0.46, p-value 0.02).

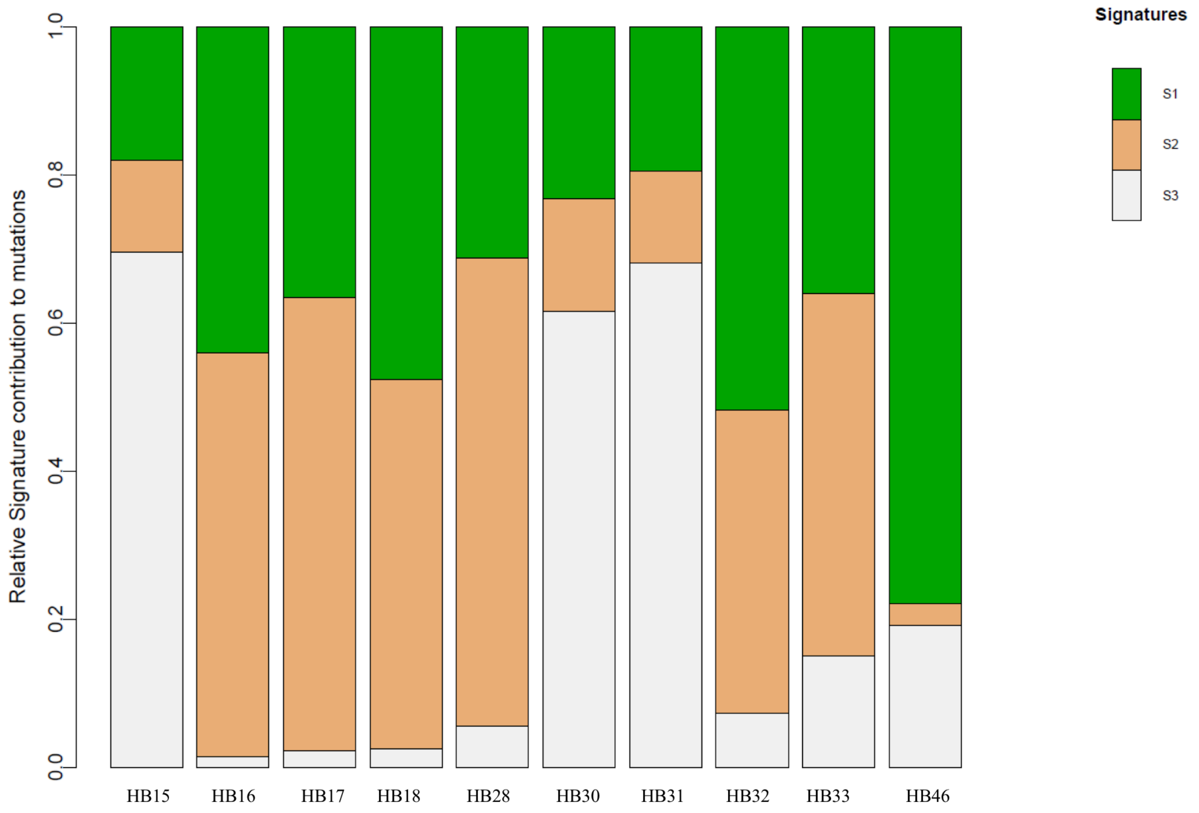


**Supplementary Figure 3:** Relative signature contribution to mutational profile of each hepatoblastoma sample.

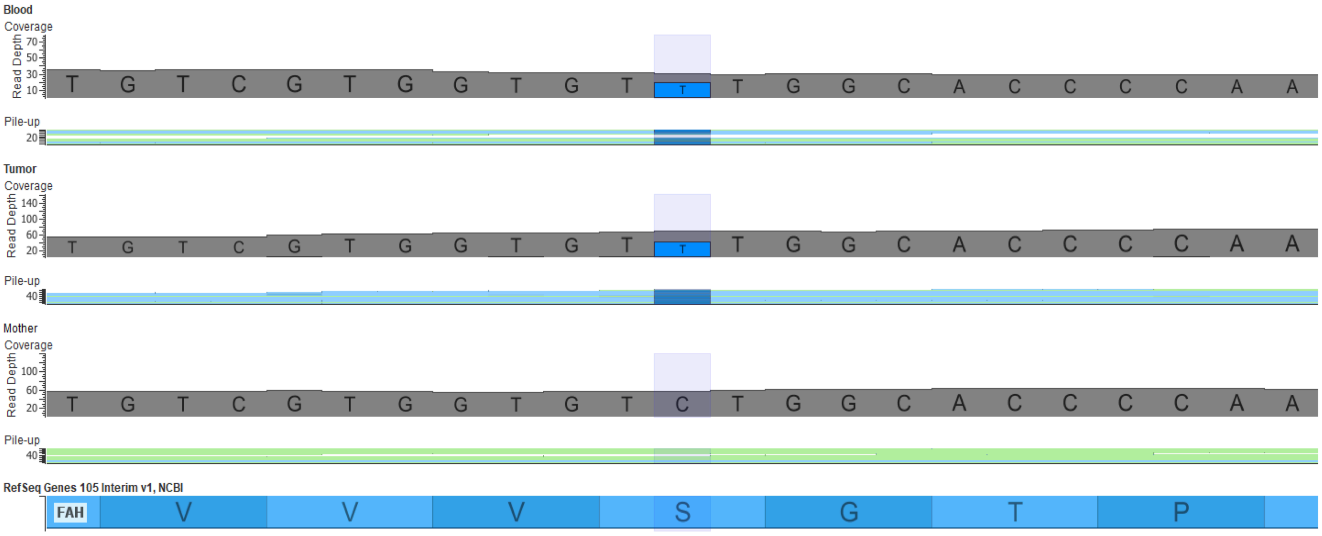


**Supplementary Figure 4: Germline *FAH* likely pathogenic variant detected by exome sequencing.** BAM file images showing a germline C>T variant (blue) in the exon 6 of the *FAH gene,* present in heterozygosity in the patient's blood and tumor, and absent from the clinically normal mother. This missense variant (p.Ser169Phe) is predicted as pathogenic by 6 out of 6 *in silico* tools, and was not previously reported in any consulted populational databases.
